# Fine modulation of carbon flow in the central carbon metabolism via ribosome-binding site modification in *Escherichia coli*

**DOI:** 10.1101/2025.08.05.668782

**Authors:** Shogo Sawada, Tatsumi Imada, Hikaru Nagai, Philip Mundt, Fumio Matsuda, Hiroshi Shimizu, Yoshihiro Toya

## Abstract

Optimization of flux distribution in central carbon metabolism is important to improve the microbial productivity. As the number of precursors required for synthesis differs for each target compound, optimal flux distribution also varies. A library of mutant strains with diverse flux distributions can aid in optimal strain screening. Therefore, in this study, we aimed to construct a library of *Escherichia coli* strains with stepwise changes in flux distribution by introducing mutations into the ribosome-binding sites of key enzyme genes on its chromosome. We focused on the flux ratios at the glucose-6-phosphate and acetyl-CoA nodes to enhance mevalonate production. Mutations were introduced into the ribosome-binding sites of *pgi* and *gltA* to vary the flux ratios of the two pathway branches. Furthermore, a combinatorial repression library comprising 16 strains was constructed by varying *pgi* and *gltA* expression at four levels, and a plasmid expressing mevalonate synthesis genes was introduced into each strain. Batch cultures were performed to obtain strains with mevalonate titers and yields were 2.4- and 3.4-fold higher than those of the parent strain. Overall, our combinatorial suppression library of *pgi* and *gltA* facilitated the effective identification of mutants with optimal metabolism for mevalonate production.

## 1. Introduction

Various useful compounds are produced using renewable carbon sources via diverse metabolic pathways in microorganisms. However, carbon flow optimization in these pathways is necessary for increased production. Central carbon metabolism plays a key role in supplying carbon backbones and energy and redox cofactors. The demand for precursor metabolites and cofactors varies depending on the target compound. For example, mevalonate, a key intermediate in the flavonoid synthesis pathway, is synthesized from three acetyl-CoA (AcCoA) molecules using two NADPH molecules. Metabolite flow optimization in pathways is necessary to satisfy the cofactor demands. Enzyme gene deletions drastically rewire the flux distribution; therefore, fine-tuning of the enzyme expression is preferable to optimize the flux distribution for enhanced target production. As metabolic pathways include multiple branches, independent alteration of flux ratios at multiple branch points is necessary to broadly change the overall flux distribution of the pathway. Therefore, a library of strains with diverse flux distributions is needed as the introduction of target-specific plasmids into suitable strains will facilitate the simple selection of strains with enhanced target production.

Gene expression level is determined by both transcriptional regulation, which depends on the promoter strength, and translational regulation, which depends on the ribosome-binding site (RBS) strength and mRNA stability. Various methods have been developed to control enzyme expression at the transcriptional and translational levels. For example, combinatorial gene repression using synthetic small regulatory RNAs enhances the production of useful compounds, such as cadaverine, phenol, and L-tyrosine (Na *et al*., 2013; Kim *et al*., 2024; Myhrvold and Silver, 2015; Pontrelli *et al*., 2018). The dynamic range is typically approximately 10-fold, limiting the control range for flux adjustment (Li *et al*., 2021). Clustered regularly interspaced palindromic repeat interference (CRISPRi) exhibits a larger dynamic range than small RNAs and is used to alter the metabolic flux in *Escherichia coli* (Donati *et al*., 2021). However, a study evaluating the proteome after suppressing the expression of 30 central metabolic genes using CRISPRi revealed that CRISPRi reduces the amount of target enzymes to an average of 0.2-fold in *E. coli*, (Lucks *et al*., 2011). Additionally, various gene suppression technologies, such as riboregulators, small transcription activating RNAs, toehold switches, and antisense RNAs, have been developed. However, diverse challenges, such as unintended gene suppression and limited dynamic range, necessitate the development of more versatile gene suppression techniques (Chappell *et al*., 2017; Green *et al*., 2014; Bonde *et al*., 2016).

In this study, we aimed to alter the flux ratio stepwise at a pathway branch point by introducing mutations into the RBSs of key enzyme genes. RBS is a sequence upstream of the start codon in the mRNA where the ribosome initially binds. RBS sequence considerably impacts the translation efficiency with even a few nucleotide changes causing significant variations in the translation levels (Oesterle *et al*., 2017). Moreover, RBS is easy to modify as its sequence length is much shorter than that of the promoter region. We selected genes encoding enzymes catalyzing the branching-point reactions in pathways as RBS modification targets (Fig. 1). Subsequently, we constructed a library of *E. coli* mutants with stepwise alterations in the flux distribution of central metabolic pathways by introducing mutations into chromosomes using multiplex automated genome engineering (MAGE). Alteration of flux ratios at the glucose-6-phosphate (G6P) and AcCoA nodes produced AcCoA derivatives by consuming NADPH for synthesis. The strain library was effective in identifying high mevalonate producers.

**Fig. 1.**
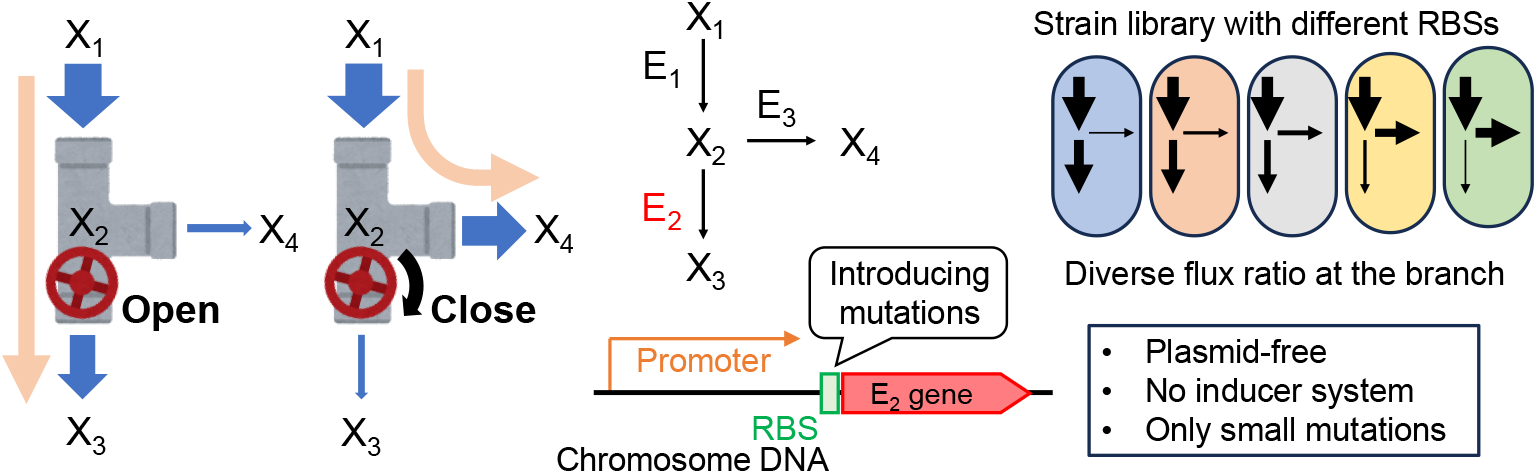
Flux ratio control at a pathway branch point by introducing mutations into the ribosome-binding site (RBS). (A) Metabolic flow was altered by repressing enzyme E2 expression. (B) Mutations were introduced into the RBS sequence of the chromosomal E2 gene. X1–X4 are metabolites, and E1–E3 are enzymes catalyzing the corresponding reactions (indicated by arrows). (C) A library of strains with diverse flux ratios at the branch point was constructed by introducing RBS mutations into the chromosome.

## 2. Materials and methods

### 2.1. Strain construction

In this study, *E. coli* MG1655(DE3) strain (Tokuyama *et al*., 2014) was used as a parental strain. MAGE oligo design tool (MODEST) was used to design the RBS sequence to repress the target enzyme expression (Bonde *et al*., 2014). Mutations introduced into the *E. coli* chromosome using the pORTMAGE-2 plasmid (Nyerges *et al*., 2016) with the primers listed in Table S1, and were confirmed via Sanger sequencing. Loss of the pORTMAGE-2 plasmid was confirmed via replica plating. Subsequently, pCOLADuet-mvaES plasmid carrying the codon-optimized *mvaE* and *mvaS* genes of *Enterococcus faecalis* was introduced into the strains for mevalonate production (Wada *et al*., 2017).

### 2.2. Culture conditions

96-well plate: Single colonies were inoculated into a 96-well plate containing 200 µL of Lennox medium and incubated overnight in Deep Well Maximizer (M·BR-022UP; TAITEC, Saitama, Japan) at 30°C with shaking at 1000 rpm. Additionally, 1 μL of culture was inoculated into a 96-well plate containing 200 µL of Wolfe’s M9 (M9W) medium, incubated at 37°C, and monitored using a microplate reader (Synergy H1 or Synergy HTX; BioTek, Winooski, VT, USA). Optical density at 600 nm (OD_600_) was measured every 30 min. L-shaped tube: A single colony was inoculated into 5 mL of Lennox medium in test tubes and incubated overnight at 37°C with shaking at 150 rpm. After washing with 5 mL of Dulbecco’s phosphate-buffered saline, the cell pellet was suspended in 2.5 mL of the same buffer. The starter was inoculated into 5 mL of M9W medium in an L-shaped tube at an initial OD_600_ of 0.05 or 0.1, and cultured at 37°C with shaking at 70 strokes per min using a Bio-Photorecorder (TVS062CA; Advantec, Tokyo, Japan). Optical density at 660 nm (OD_660_) was measured every 5 min. Shake flask scale: A single colony was inoculated into 5 mL of Lennox medium in test tubes and incubated overnight at 37°C with shaking at 150 rpm. After washing with 5 mL of M9W medium, the cell pellet was suspended in 2.5 mL of same medium. The starter was inoculated into 50 mL of M9W medium in a 200-mL baffled flask at an initial OD_600_ of 0.05, and cultured at 37°C with 150 rpm. For mevalonate production, 100 µM isopropyl β‐D‐1‐ thiogalactopyranoside (IPTG) was added to the broth (Wada *et al*., 2017).

### 2.3. Citrate synthase assay

The *E. coli* cells were cultured in flask-scale conditions and collected from 10 mL of broth via centrifugation at 8000 × g for 5 min at 4°C when the OD_600_ reached 0.6–0.8. After washing with 5 mL of M9W medium, the cell pellet was stored at -80°C. The cells were suspended in 500 µL of lysis buffer (50 mM Tris-HCl [pH 8.1] and 150 mM NaCl) containing the ProteoGuard EDTA-free protease inhibitor cocktail (Clontech, Mountain View, CA, USA) and sonicated using Bioruptor (UCD-205; Diagenode, NJ, USA) for ten cycles (30 s on/30 s off at high power). The crude cell extract was collected via centrifugation at 18000 × g for 20 min at 4°C, and diluted in the same buffer to 0.5 mg mL^-1^. Protein concentration was measured using a microvolume spectrophotometer µLite+ (BioDrop, Cambridge, UK). GltA catalyzes the citrate synthase reaction converting AcCoA and oxaloacetate (OAA) to citrate and reduced CoA, which reacts with 5,5’-dithiobis(2-nitrobenzoic acid) (DTNB) to form 5-thio-2-nitrobenzoic acid that exhibits absorbance at 412 nm (Srere *et al*., 1969). The assay solution was prepared by mixing DTNB (100 µL, 1 mM), AcCoA (30 µL, 10 mM), and Milli-Q water (770 µL). After adding 850 µL of the assay solution to a cuvette with 100 µL of the crude enzyme extract, absorbance at 412 nm was recorded for 3 min to determine the AcCoA deacylase activity. Then, 50 µL of 10 mM OAA was added to the cuvette, and absorbance at 412 nm was recorded for 3 min to determine the citrate synthase activity.

### 2.4. Flux ratio analysis

The cells were cultured using [1-^13^C] glucose as the sole carbon source. During the exponential phase, the cultures (2 mL) were harvested and centrifuged at 10000 × g for 5 min at 4°C. The cell pellet was washed twice with 0.9% NaCl and hydrolyzed using 2 mL of 6 M HCl at 105°C for 18 h. After filtration and evaporation, proteinogenic amino acids were derivatized via *tert*-butyl-dimethylsilylation. Mass isotopomers of alanine were measured using a gas chromatography-mass spectrometer (Shimadzu, Kyoto, Japan) equipped with the DB-5MS column (Agilent Technologies, Santa Clara, CA, USA). The flux ratio among the glycolytic pathways was calculated from the mass isotopomer distribution of the [M-57]^+^ and [M-85]^+^ fragment ions of proteinogenic alanine. All analytical and calculation procedures have been previously described (Morita *et al*., 2017).

### 2.5. High-performance liquid chromatography analysis

Culture supernatants were obtained via centrifugation at 20400 g for 5 min at 4°C. Extracellular glucose, acetate, and mevalonate concentrations were quantified using a high-performance liquid chromatography system (Shimadzu) equipped with the Aminex HPX-87H column (Bio-Rad, Hercules, CA, USA), as previously described (Miyoshi *et al*., 2023).

## 3. Results

### 3.1. G6P isomerase (PGI) repression library construction

*E. coli* has three glycolytic pathways: Embden–Meyerhof–Parnas (EMP), pentose phosphate (PP), and Entner–Doudoroff (ED) pathways (Fig. 2A). PGI catalyzes the conversion of G6P to fructose-6-phosphate, and its expression alters the flux ratio at the G6P node (Kamata *et al*., 2019). To downregulate PGI expression, mutations in the RBS of *pgi* were predicted using MODEST (Bonde *et al*., 2014). We previously reported that significant reduction of PGI expression is necessary to alter the flux (Tander *et al*., 2019). Therefore, we designed mutated RBSs with expression ratios of 0.0100, 0.0053, and 0.0014 compared to the wild-type (WT; RBS a–c; Fig. 2B). Using oligonucleotide DNA with the predicted mutations (Table S1), PGI a–c strains were constructed from the MG1655(DE3) strain via MAGE. By introducing artificial DNA oligonucleotides functioning as Okazaki fragments into cells with inhibited mismatch repair mechanisms, the mutations were incorporated into arbitrary sequences on the chromosome.

**Fig. 2.**
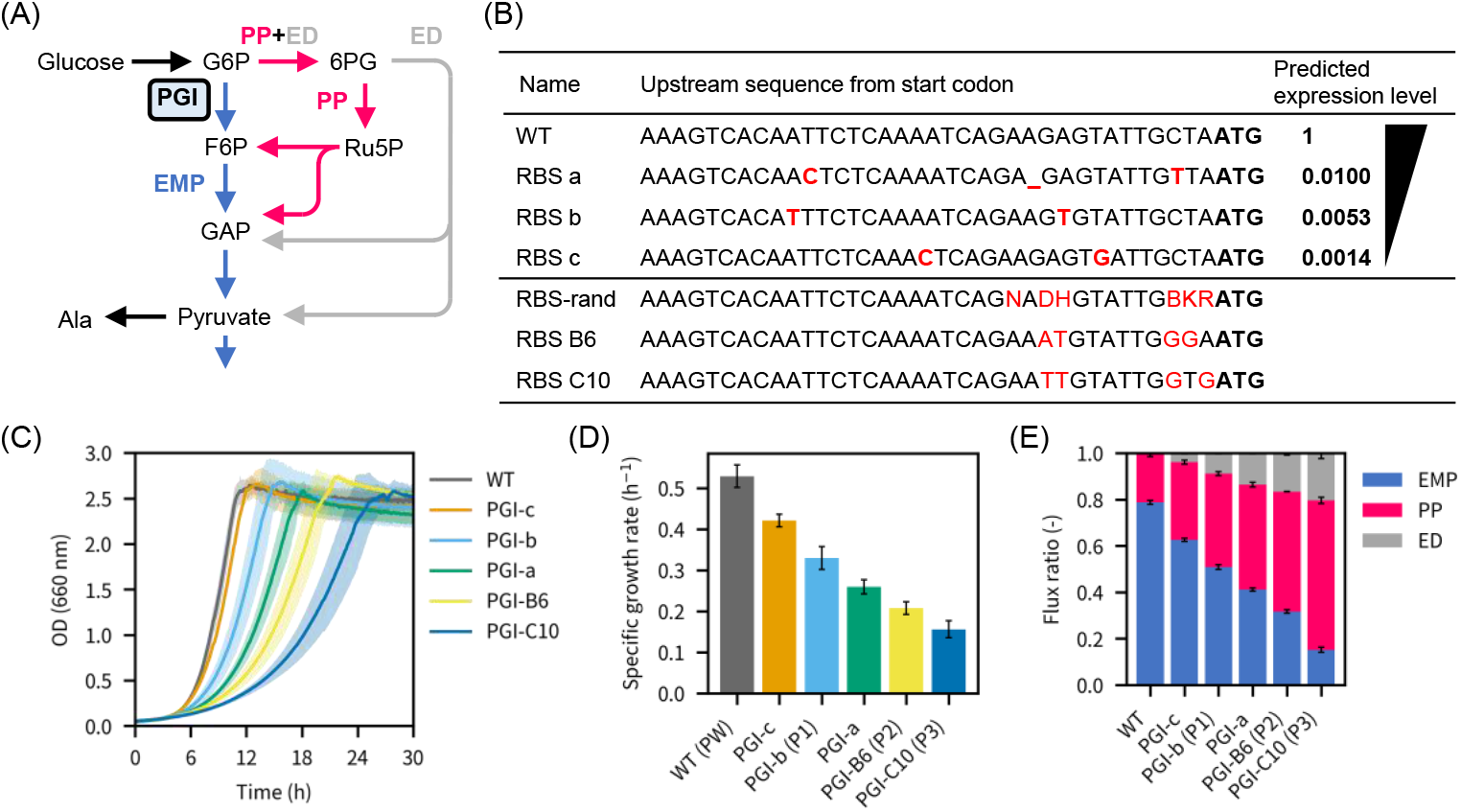
Glucose-6-phosphate isomerase (PGI) RBS engineering to alter the flux ratio at the G6P node. (A)*E. coli* glycolytic pathways. Flux ratio between the Embden–Meyerhof–Parnas (EMP) and pentose phosphate (PP) pathways was altered by repressing PGI expression. (B)Upstream sequences of *pgi* in the wild-type (WT) and engineered strains. Mutations are indicated in red. (C) Growth profile. Shaded regions indicate the standard deviations of triplicate experiments. (D) Specific growth rates. (E) Flux ratios among the EMP, PP, and Entner–Doudoroff (ED) pathways. Error bars represent the standard deviations of triplicate experiments.

When the glycolytic pathway used for glucose catabolism shifts from the EMP pathway to PP pathway, the growth rate decreases. PGI a–c and WT strains were cultured in a minimal glucose medium. Although the order of the predicted PGI expression levels and growth rates differed, growth rates of PGI a–c strains showed a stepwise decrease (Fig. 2C and D). To determine the flux ratio between the EMP and PP pathways in these strains, the cells were cultured with [1-^13^C] glucose. Flux ratio was calculated using mass isotopomers of proteinogenic alanine (Fig. 2E). Indeed, high flux ratio in the EMP pathway led to a high growth rate.

Even in the PGI-a strain, which exhibited the lowest EMP pathway flux ratio, the value was 0.38 (Fig. 2E). Therefore, we attempted to generate mutants with stronger PGI inhibition. PGI a–c strain results suggest that, although MODEST predicts the expression-suppressing mutations, it does not accurately predict the expression levels. Therefore, we used MODEST to output multiple candidate mutations and subsequently generated mutants using oligo-DNA-containing random sequences that included these mutations. As the EMP flux ratio is associated with the growth rate, strains with low EMP flux ratios were screened using growth as an indicator. We cultured the generated mutants in 96-well plates and selected two slow-growing strains (PGI-B6 and PGI-C10). Growth characteristics of the PGI-B6 and PGI-C10 strains in L-shaped tubes are shown in Fig. 2C and D. Specific growth rates of both mutants were intermediate between those of the PGI-a and Δ*pgi* strains. Furthermore, EMP flux ratios of the PGI-B6 and PGI-C10 strains were lower than that of the PGI-a strain. RBS sequencing revealed that each gene contained four base mutations (Fig. 2B). A library was successfully constructed in which the flux gradually shifted from the EMP to PP pathway.

### 3.2. GltA repression library construction

Next, we attempted to alter the flux ratio between the tricarboxylic acid (TCA) cycle and acetate overflow by repressing GltA expression (Fig. 3A). GltA catalyzes the conversion of AcCoA and OAA to citrate, and its expression alters the flux ratio at the AcCoA node. As some essential building blocks, including glutamate, are synthesized from α-ketoglutarate (αKG) via the TCA cycle with glucose as the carbon source, GltA repression mutants were screened using growth as an indicator. A mutated RBS with an expression ratio of 0.01 relative to WT was designed using MODEST (RBS 0.01; Fig. 3B) to generate the GltA-0.01 strain. Additionally, we generated 27 candidate mutated RBS sequences with gradually reduced expression levels as well as oligo-DNA-containing mutations frequently found in the resulting sequences (Fig. 3B). We selected small colonies, cultured the mutants in a 96-well plate, and selected two slow-growing strains (GltA-F4 and GltA-C5). Growth profiles and specific growth rates of the WT, GltA-0.01, GltA-F4, and GltA-C5 strains in L-shaped tubes are shown in Fig. 3C and D, respectively. RBS sequencing revealed that GltA-F4 and GltA-C5 contained two base mutations (Fig. 3B). GltA activity indicated a stepwise change in GltA expression (Fig. 3E).

**Fig. 3.**
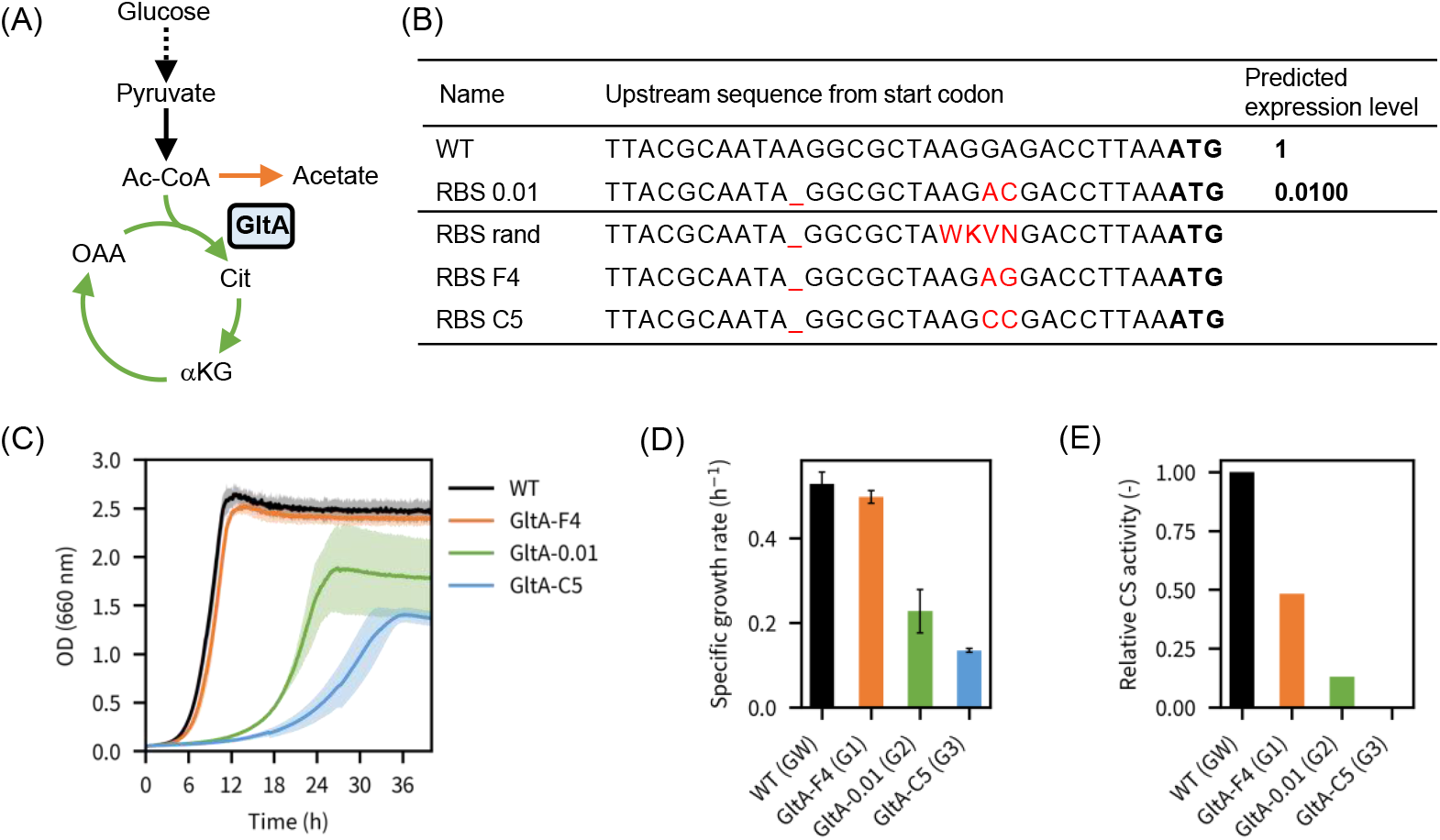
GltA RBS engineering to alter the flux ratio at the acetyl-CoA (AcCoA) node. (A) *E. coli* metabolic pathways around the AcCoA node. Flux ratio between the tricarboxylic acid (TCA) cycle and acetate overflow was altered by repressing GltA expression. (B) Upstream sequences of *gltA* in the WT and engineered strains. Mutations are indicated in red. (C) Growth profiles. Shaded regions indicate the standard deviations of triplicate experiments. (D) Specific growth rate. Error bars represent the standard deviations of triplicate experiments. (E) GltA activity.

### 3.3. Combinatorial repression of PGI and GltA

Flux ratio at the G6P node affects the NADPH supply, whereas that at the AcCoA node affects the AcCoA availability (Kamata *et al*., 2019). As some useful compounds, including mevalonate, are synthesized from AcCoA using NADPH as a redox cofactor, we performed PGI and GltA combinatorial repression. Using oligo DNAs for PGI repression (Fig. 2B), WT and GltA repression strains were transformed via MAGE.

For the PGI inhibition series, RBSs of WT, RBS-b, RBS-B6, and RBS-C10 were selected from seven steps (Fig. 2B and E) and labeled as PW, P1, P2, and P3, respectively. Similarly, for the GltA inhibition series, RBSs of WT, RBS-F4, RBS-0.01, and RBS-C5 were labeled as GW, G1, G2, and G3, respectively (Fig. 3B and E). In total, 16 strains were constructed by altering the PGI and GltA expression at four different levels, and these strains were cultured in L-shaped tubes (Fig. 4). Synergistic growth reduction was observed upon PGI and GltA suppression (Fig. 4B). Glycolytic flux ratios of PWGW, P1G1, P2G2, P3GW, and PWG3 were analyzed using [1-^13^C] glucose (Fig. 4C). Notably, EMP flux ratio decreased stepwise with decreased PGI suppression in the PWGW, P1G1, and P2G2 strains. Flux ratio at the G6P node reflected the PGI suppression level in the P3GW and PWG3 strains. Specific glucose consumption and acetate production rates are shown in Fig. 4D. Acetate overflow proportion increased with increased GltA inhibition.

**Fig. 4.**
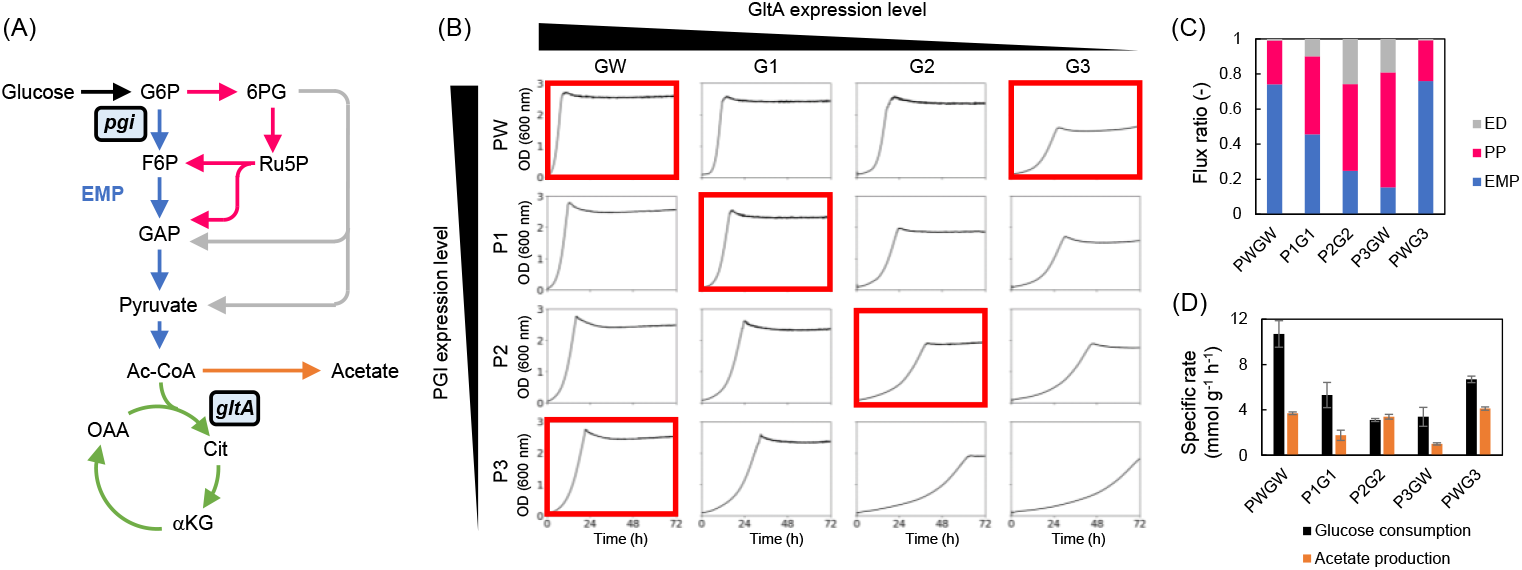
Culture profiles of the PGI and GltA combinatorial repression mutants. (A) PGI- and GltA-catalyzed reactions in central carbon metabolism. (B) Growth profiles. (C) Flux ratios. (D) Specific rates.

### 3.4. Mevalonate production using the combinatorial mutation library

As mevalonate is synthesized from three AcCoA and two NADPH molecules, regulation of AcCoA and NADPH supply is important to increase the mevalonate yield (Fig. 5A). PGI and GltA inhibition is an effective approach to accumulate these precursors; however, these enzyme deficiencies can have a fatal impact on growth. Therefore, mevalonate productivity was evaluated in terms of yield and production rate, using stepwise PGI- and GltA-suppressed mutants.

**Fig. 5.**
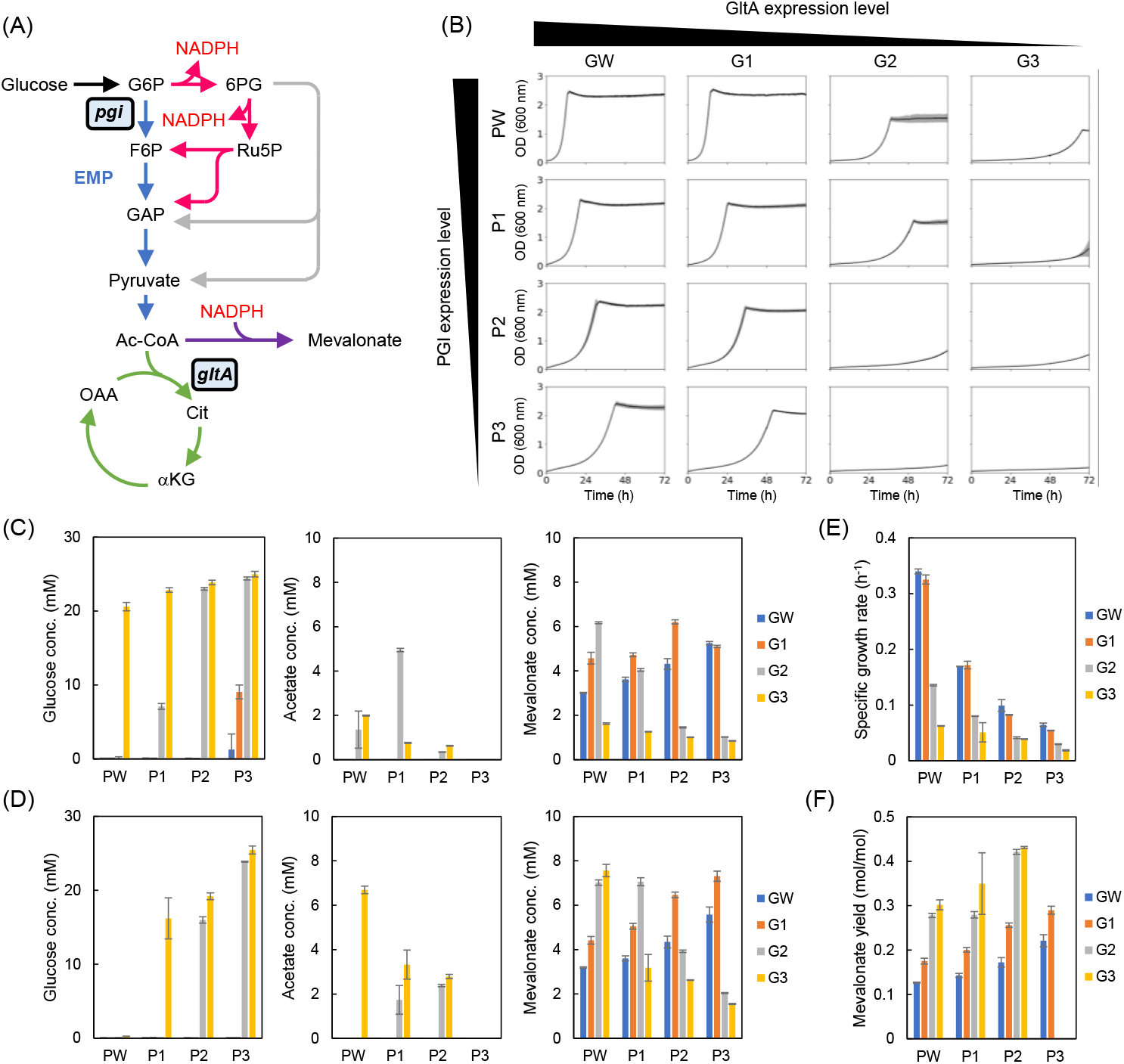
Mevalonate production in the PGI and GltA combinatorial repression mutants. (A) Mevalonate synthesis pathway and central carbon metabolism. (B) Growth profiles. Shaded regions indicate the standard deviations of triplicate experiments. Extracellular metabolite concentrations at (C) 48 h and (D) 72 h. (E) Specific growth rates during the exponential phase. (E) Mevalonate yield after 72 h. Mevalonate yields of P3G2 and P3G3 could not be calculated due to low glucose consumption. Error bars represent the standard deviation of triplicate experiments.

A plasmid expressing MvaE and MvaS, which catalyze the conversion of AcCoA to mevalonate, was introduced into 16 combinatorial repression strains. Growth profiles of these strains are shown in Fig. 5B. The decreased growth rate tendency due to PGI and GltA suppression is similar to that shown in Fig. 4C, and an overall decrease in growth rate was observed after the plasmid introduction. Notably, strains with strong repression (P1G3, P2G2, P2G3, P3G2, and P3G3) did not reach full growth even after three days of culture. Extracellular glucose and mevalonate concentrations at 48 and 72 h are shown in Fig. 5C and D, respectively. Specific growth rates during the exponential phase and overall mevalonate yields are shown in Fig. 5E and F, respectively.

After 48-h culture, PWG2 and P2G1 strains produced over 6 mM mevalonate (Fig. 5C). Highest production was achieved in combination with P3 in the GW-series strains and P2 in the G1-series strains. In the G2- and G3-series strains, highest production was achieved in combination with PW.

After 72-h culture, PWG3, P1G2, and P3G1 strains produced over 6 mM mevalonate (Fig. 5D). Highest titer was observed in the PWG3 strain, which was 2.4-fold higher than that in the PWGW strain. PWG3 strain retained glucose and exhibited the potential to produce mevalonate under extended culture conditions. Mevalonate yield was high in the P2G2 and P2G3 strains, exceeding 0.4 mol mol^-1^. Notably, yield increased with increased PGI suppression. This suggests that mevalonate yields of the P3G2 and P3G3 strains are possibly even higher; however, yields of these strains could not be accurately evaluated due to their low glucose consumption.

## 4. Discussion

Engineering methods altering the flux distribution in central carbon metabolism are highly desirable to improve useful compound production (Kim *et al*., 2018; Kwak *et al*., 2020; Dodelin *et al*., 2025). In this study, we developed a method to achieve stepwise changes in the flux ratio by introducing mutations into the RBSs of key enzyme genes on the chromosome. Using this method, we constructed an *E. coli* strain library with diverse flux ratios at the G6P and AcCoA nodes. Focusing on enzymes catalyzing reactions with high flux at the pathway branch, flux ratios were altered by suppressing their expression. Uncertainty in predicting the expression strength of the RBS sequence was addressed by associating growth with target enzyme activity for easy candidate screening. This method effectively altered the overall flux distribution of the pathways by combining mutations at multiple branch points.

In this study, a combinatorial repression library comprising 16 strains was constructed by varying PGI and GltA expression at four levels. Subsequently, mevalonate production was assessed using this strain library. Compared to the parental PWGW strain, some mutants showed increased yield and production rates (up to 2.4- and 3.4-fold higher, respectively). At all GltA suppression levels, high PGI suppression tended to result in a high mevalonate yield. Notably, a trade-off between yield and production rate was observed during mevalonate production. Specifically, yield increased but production rate decreased with increased PGI and GltA inhibition. As mevalonate production was evaluated in batch cultures, cell growth decreased under extremely high PGI and GltA suppression, resulting in low mevalonate titers. These results suggest that the optimal strain differs depending on whether the production rate or yield is. PWG2 and P2G1 strains are preferable if the production rate is prioritized, whereas the P2G2 and P2G3 strains are better choices if the yield is prioritized. Under extended culture times exceeding 72 h, P3 series strains are preferable for high mevalonate yield. This trend is consistent with our previous findings that strong PGI inhibition is necessary to achieve a high mevalonate yield during the growth phase (Kamata *et al*., 2019). Another study reported the effects of flux manipulation of the upstream glycolysis pathway and TCA cycle on mevalonate production in *E. coli*. Dodelin *et al*. (2025) introduced amino acid substitution mutations to decrease the PfkA and GltA activities and evaluated their impacts on mevalonate production. Mevalonate yield increased as GltA activity decreased, consistent with our results. Whether introducing mutations to reduce the enzyme activity or introducing mutations into RBS to suppress enzyme expression is a better strategy to decrease the flux is an important consideration. Directly introducing mutations into the enzyme can alter its function; however, introducing mutations into RBS does not alter the enzyme function. Moreover, quantitative prediction of the ways in which amino acid substitutions affect enzyme activity remain difficult. Therefore, introducing mutations into the RBS is a promising strategy to change the pathway flux ratio.

Importantly, introduction of unnecessary plasmids is not necessary to modify the RBS sequence on the chromosome. We previously evaluated mevalonate production by adjusting the flux ratio at the G6P node using an IPTG-induction system for precise expression control (Kamata *et al*., 2019). However, in this study, IPTG-inducible system could not be used to express the target synthesis enzymes. Generally, synthesis pathway enzymes should be highly expressed, and excellent inducible systems, such as IPTG and arabinose-inducible promoters, are not desirable for fine-tuning the flux. Furthermore, types of available induction chemicals are limited when multiple branching points are independently controlled. Therefore, for cost management, these chemicals should not be used to produce inexpensive targets. Notably, our approach facilitates the manipulation of metabolic fluxes without using inducible systems. Overall, our findings highlight the importance of mutant libraries with broad flux distributions for the identification of host strains with optimal metabolic states for efficient target production.

## CRediT authorship contribution statement

**Shogo Sawada:** Validation, Investigation. **Tatsumi Imada:** Validation. **Hikaru Nagai:** Investigation. **Philip Mundt:** Validation, **Fumio Matsuda:** Methodology, Supervision. **Hiroshi Shimizu:** Conceptualization, Writing – review & editing, Supervision. **Yoshihiro Toya:** Conceptualization, Methodology, Investigation, Validation, Writing – original draft.

## Acknowledgments

pORTMAGE-2 was a gift from Csaba Pál (Addgene, plasmid # 72677).

## Data available

Data will be made available on request.

---

**Table S1.**
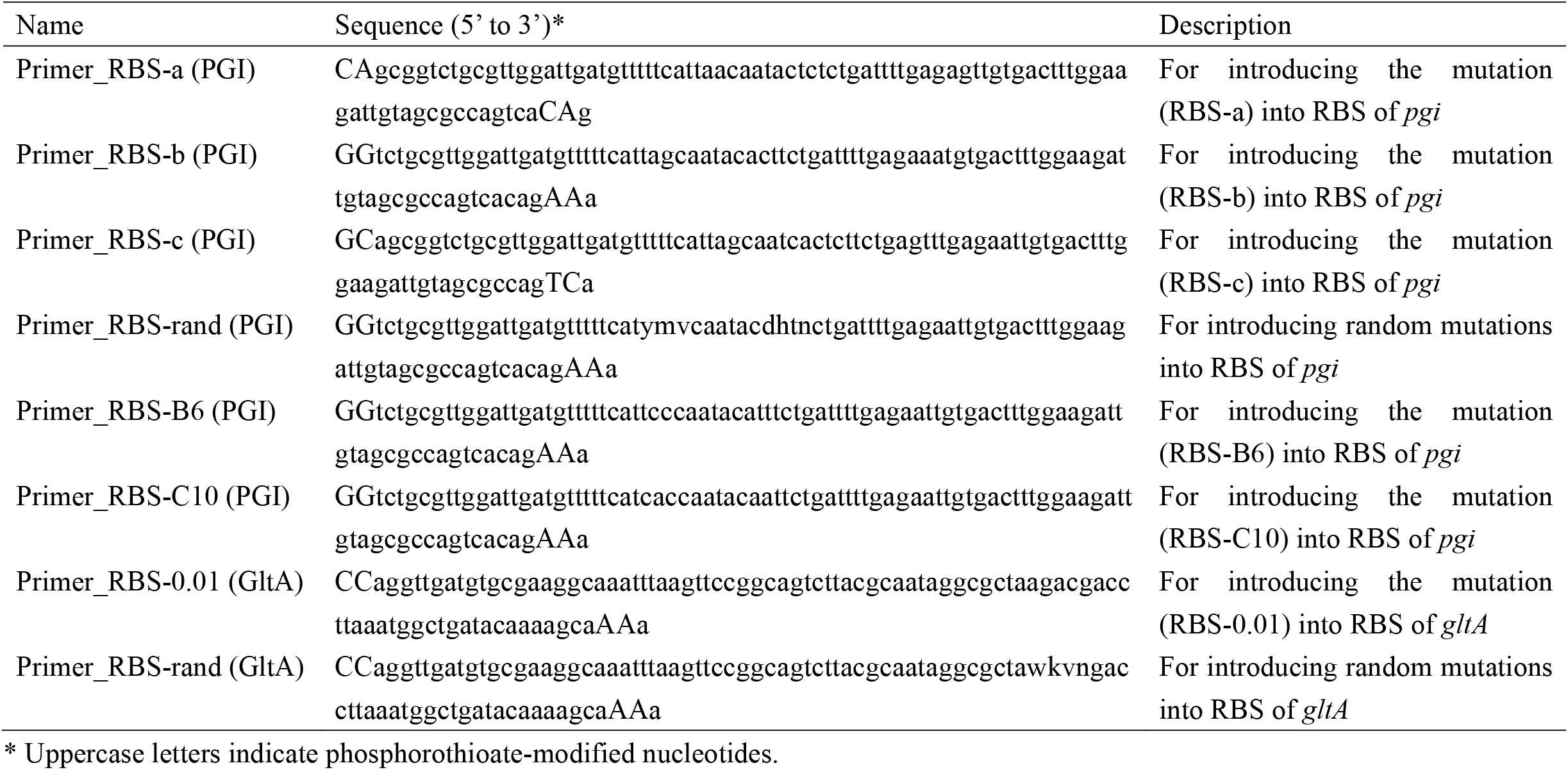
Primer list for introducing mutations by MAGE.

